# History information emerges in the cortex during learning

**DOI:** 10.1101/2022.11.01.514667

**Authors:** Odeya Marmor, Fritjof Helmchen, Ariel Gilad

## Abstract

We learn from our experience but the underlying neuronal mechanisms incorporating past information to facilitate learning is relatively unknown. Specifically, which cortical areas encode history-related information and how is this information modulated across learning? To study the relationship between history and learning, we continuously imaged cortex-wide calcium dynamics as mice learn to use their whiskers to discriminate between two different textures. We mainly focused on comparing the same trial type with different history information, i.e., a different preceding trial. We found history information in barrel cortex (BC) during stimulus presentation. Importantly, history information in BC emerged only as the mouse learned the task. Next, we also found learning-dependent history information in rostrolateral (RL) association cortex that emerges before stimulus presentation, preceding activity in BC. History information was also found in other cortical areas and was not related to differences in body movements. Interestingly, a binary classifier could discriminate history information at the single trial level just as well as current information both in BC and RL. These findings suggest that past experience emerges in the cortex around the time of learning, starting from higher-order association area RL and propagating down (i.e., top-down projection) to lower-order BC where it can be integrated with incoming sensory information. This integration between the past and present may facilitate learning.

## Introduction

Learning is a process of acquiring new knowledge required for appropriate behavior and is highly dependent on our previous experience. Our brain integrates incoming sensory information with history information of previous stimuli to form a knowledgeable association of the current stimulus. Despite the strong link between history and learning, the underlying cortex-wide dynamics are relatively unknown, partially because most previous studies separately focus either on learning or history(Hattori et al., 2019). Learning-related neuronal dynamics are broadly observed across the whole cortex, including primary sensory or motor areas(Blake et al., 2002; Chen et al., 2015; Gilad and Helmchen, 2020; Jurjut et al., 2017; Komiyama et al., 2010; Li et al., 2008; Poort et al., 2015; Wiest et al., 2010; Xu et al., 2014; Yan et al., 2014), higher-order association areas(Driscoll et al., 2017b; Gilad and Helmchen, 2020) and prefrontal cortex(le Merre et al., 2018; Pasupathy and Miller, 2005). But do these areas that participate in the learning process also carry history information?

Encoding of history information has been reported mainly in higher order cortical areas such as the posterior parietal cortex (PPC)(Akrami et al., 2018; Harvey et al., 2012; Hwang et al., 2017; Morcos and Harvey, 2016; Benjamin B Scott et al., 2017; Suzuki et al., 2022), retrosplenial cortex(Hattori et al., 2019; Vann et al., 2009) and prefrontal cortex (Banerjee et al., 2020; Johnson et al., 2016; Kawai et al., 2015; Benjamin B. Scott et al., 2017; Sul et al., 2010; Tsutsui et al., 2016), but also to a smaller extent in lower-order primary sensory areas such as BC(Banerjee et al., 2020; Chéreau et al., 2020; Rodgers et al., 2021). There is still a debate on which areas link history information with the learning process. Another important aspect of the history-learning relationship is the temporal aspect that enables integration of past information with present sensory information. For example, does history information emerges in cortex before present information arrives or do both past and present information maybe emerge simultaneously in a certain cortical area? From the temporal aspect, optogenetic silencing of PPC area during the inter-trial interval affected performance, highlighting that higher-order cortical areas may maintain history information before the incoming current stimulus(Akrami et al., 2018; Hwang et al., 2017).

To study the history-learning relationship, we use wide-field cortical imaging of mice learning to discriminate between two textures and focus on the cortex-wide dynamics of history information. In a previous study using the same dataset, we showed that in mice learning a whisker-based texture discrimination task, the activity in task-related areas (e.g., barrel cortex – BC and rostrolateral association cortex – RL) increases as they become experts(Gilad and Helmchen, 2020). RL is part of the PPC and is located within the cluster of higher-order association areas surrounding V1. RL plays pivotal roles in cross-modal sensory integration, learning and history, but the relationship between history and learning in RL is unknown (Akrami et al., 2018; Driscoll et al., 2017a; Hattori et al., 2019; Hwang et al., 2017; Khodagholy et al., n.d.; Marcos and Harvey, 2016; Save and Poucet, 2009). Here, by classifying trials according to the preceding trial, we now demonstrate the emergence of history information as the mouse gains expertise. Specifically, history information emerges in RL, just before the stimulus presentation during the trials, and then is transferred to BC during the texture touch period, which may aid in learning the rewarded stimulus.

## Results

In this study we investigate history-dependent dynamics across the whole dorsal cortex and its emergence during learning in transgenic mice expressing a calcium indicator (GCaMP6f) in L2/3 excitatory neurons (n=7 mice). This dataset is identical to the one published in Gilad and Helmchen(Gilad and Helmchen, 2020) where we focused only on learning dynamics. Using wide-field calcium imaging through the intact skull (Gallero-Salas et al., 2021; Gilad et al., 2018b; Gilad and Helmchen, 2020; Vanni and Murphy, 2014), we chronically measured large-scale neocortical L2/3 activity in the contralateral hemisphere as mice learned a go/no-go whisker-dependent texture discrimination task (Gilad and Helmchen, 2020). Whisker movements and body movements were video monitored and synchronized to the calcium imaging data (Methods). To delineate areas in the dorsal cortex, we functionally mapped sensory areas for each mouse during anesthesia (see Methods). Based on these maps (and skull coordinates) we registered all images to the 2D top-view Allen reference atlas(Oh et al., 2014) and defined 25 areas of interest, further divided into four groups (Fig. 1c;(Gilad and Helmchen, 2020)).

**Figure 1.**
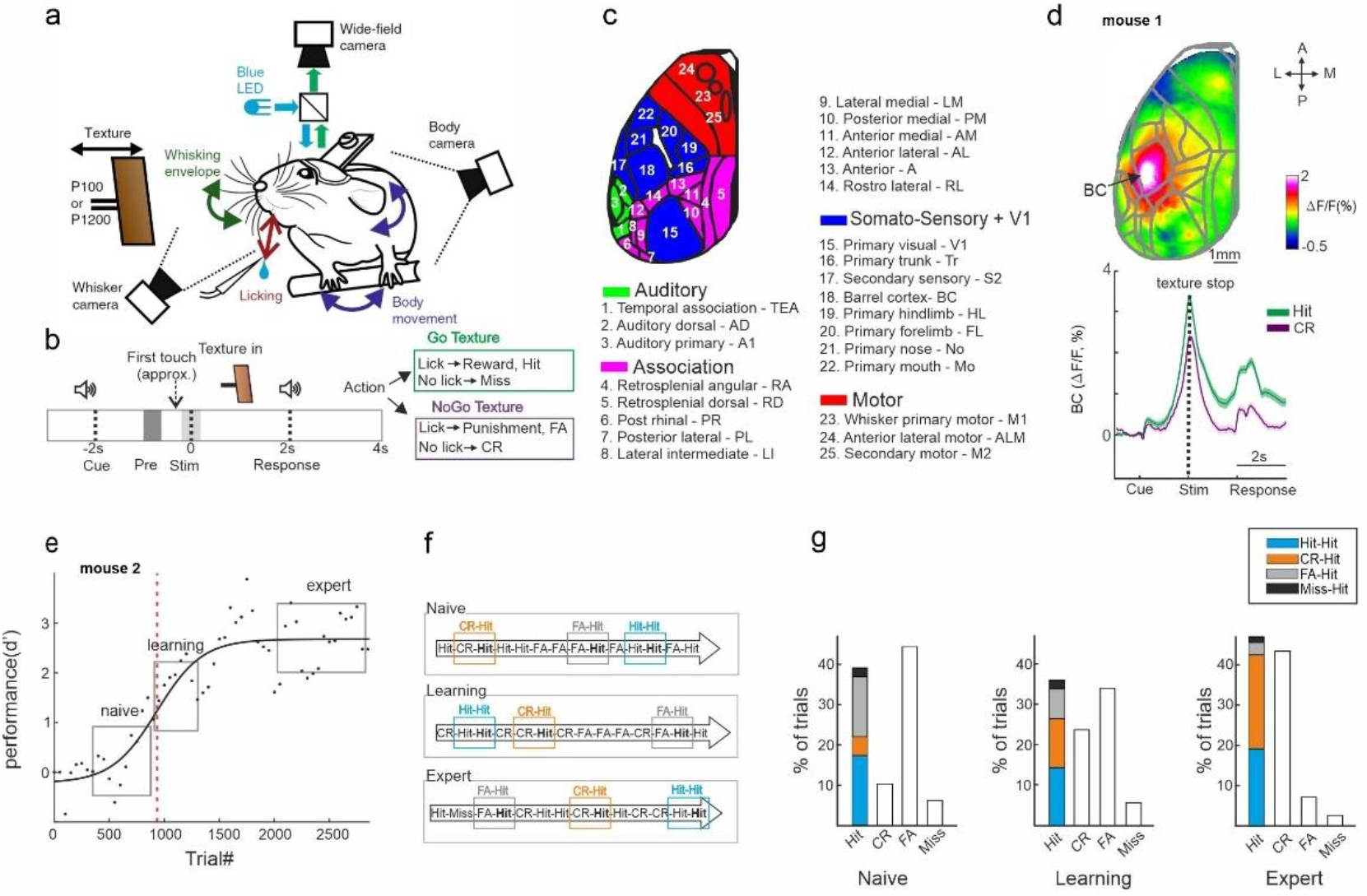
Trial types based on history. **a**. Behavioral setup for head-fixed texture discrimination with simultaneous wide-field calcium imaging and video monitoring of whisker motion and body movement. **b**. Trial structure and possible trial outcomes. pre- and stim-periods are marked in gray and light gray colors, respectively. **c**. 25 cortical areas used in this study grouped into auditory areas (green), association areas (pink), somatosensory + V1 areas (blue), and motor areas (red) **d**. *Top*: Example mean activation map (averaged during the stim period) for the Hit condition. BC – barrel cortex. Color denotes normalized fluorescence. Bottom: Time course of activity in BC for Hit (green) and CR (purple). Error bars are mean±SEM across trials (n=376 and 333 for Hit and CR respectively). **e**. Example of a learning curve (d′ as a function of trial number) of one mouse, fitted with a sigmoid function (solid black line). Red dashed vertical line indicates the learning threshold. gray rectangles mark the naive, learning and expert phases. **f**. Schematic diagram of the different trial types for a Hit trial preceded by a different trial (i.e., history): Hit-Hit (blue), CR-Hit (orange) and FA-Hit (gray). **g**. Probability of the different trial types along with the distribution of history for the Hit trial during the naïve, learning and expert phases (averaged across 7 mice).

Mice were trained on a head-fixed, whisker-based go/no-go texture discrimination task (Chen et al., 2013; Gilad and Helmchen, 2020)(Fig. 1a; Methods). Each trial started with an auditory cue (stimulus cue), signaling the approach of either two types of sandpapers (grit size P100: rough texture; P1200: smooth texture; 3M) to the mouse’s whiskers as ‘go’ or ‘no-go’ textures. The texture stayed in touch with the whiskers for 2 s, and then it was moved out after which an additional auditory cue (response cue) signaled the start of a 2-s response period (Fig. 1b) followed by a 6-s break until the next trial auditory cue. Five mice were trained to lick for the P100 and two mice were trained to lick for the P1200 texture. Mice were rewarded in ‘Hit’ trials for correctly licking after the go texture and punished with white noise for incorrectly licking for the no-go texture (‘false alarm’ trials, FA). Mice were neither rewarded nor punished when they withheld licking for the go and no-go textures (‘Miss’ and ‘correct-rejection’, CR, trials, respectively). We defined two time windows within the trial structure: the ‘pre-period’ when the texture approaches the whiskers (−1 to −0.6 s relative to the texture stop; mainly before the first whisker-texture touch); and the ‘stim-period’ during texture touch (−0.2 to 0.2 s relative to texture stop; Fig. 1b).

The performance of all mice increased with training (5–11 days; ~500 trials/day) and eventually reached high discrimination levels (quantified by d-prime; d’; Fig. s1; refs. (Gilad et al., 2018a); Methods). We defined the ‘learning threshold’ of reaching expert level for each mouse by crossing the inflection point of the sigmoid fit for the learning curve (in units of ‘trial number’; Fig. 1e, Fig. s1). The fastest learning mouse reached threshold in slightly less than thousand trials whereas mouse #4 took substantially longer (Fig. s1). In addition, we defined a naïve (1^st^ day of recording), learning (day of crossing the learning threshold; 2^nd^ or 3^rd^ day) and expert (last recording day) phases for each mouse. All mice, after gaining expertise, showed strong activation in the Barrel cortex (Fig 1. d, upper panel). This activation was during stimulus representation, stronger in Hit trials compared to CR trials (Fig. 1d, lower panel), not dependent on the texture type (i.e. if the hit was p100 or p1200).

Here, we focus on the history content for each trial type. We sub-grouped all the Hit trials (i.e., the current trial type) based on the previous trial type: CR (“CR-Hit"; n=423±74, mean±SEM), Hit (“Hit-Hit"; n=585±42), FA (“FA-Hit"; n=217±24) or Miss (“Miss-Hit"; n=55±24; Fig.1f, g). “Miss-Hit” were not analyzed due to a small number of trials. Our main analysis will compare “CR-Hit” (orange) and “Hit-Hit” (blue) trial pairs, since they are present in large numbers during all phases in each mouse separately (Fig. 1g; But see Fig. s3 for a comparison of other trial pairs). We emphasize that in this comparison, the current trial type is identical (i.e., Hit) whereas only the pervious trial (i.e., the history, CR or Hit) differed, therefore eliminating activity differences due to the current stimulus.

### History information in BC emerges during learning

First, we focused on history-dependent information in BC, specifically during the stim-period. BC displayed higher activity during CR-Hit compared to Hit-Hit only during learning and expert phases, but not during the naïve period (Fig. 2a, Fig. s2). This difference was significant during the stim-period in learning and expert phases across mice (Fig. 2b; signed rank test, p<0.05). To check whether this effect is not due to difference in body or whisker movements between the two pair types, we analyzed body movements by calculating (1 - frame-to-frame correlation) in mouth, forelimb and hindlimb areas and computed whisker envelope as a function of time (see Methods). Both body movements and whisker envelope were similar between CR-Hit and Hit-Hit pairs (Fig. 2c) and there was no significant difference across mice during the stim-period for neither naïve, learning or expert phases (Fig. 2d p>0.05; Signed rank test) nor during the pre-period (p>0.05, signed rank test, data not shown). This result, along with the fact that the current trial type in both conditions is identical, strongly indicates the presence of history information in BC.

**Figure 2.**
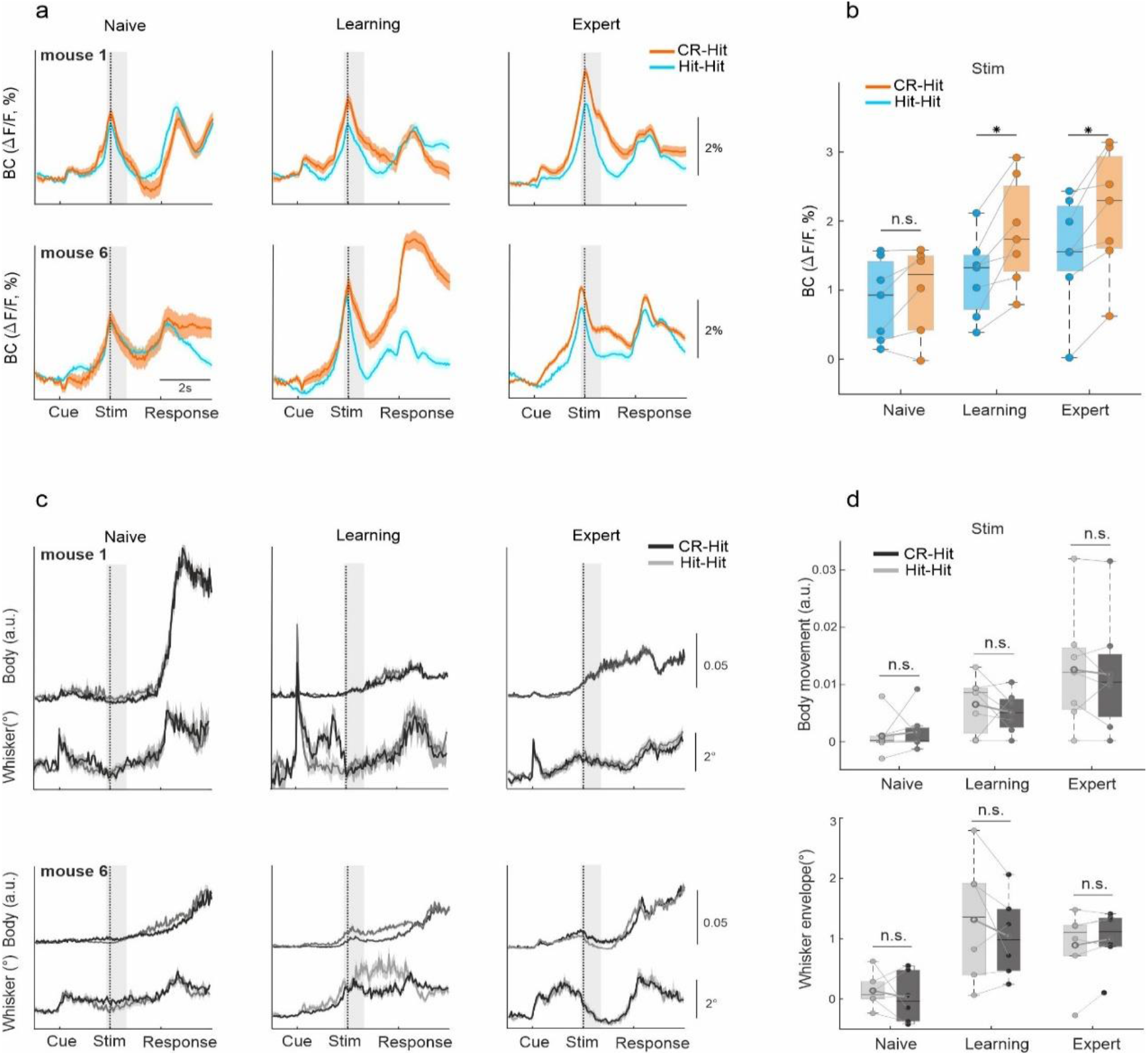
History information in BC. **a**. Example of average BC response of Hit-Hit (blue) and CR-Hit (orange) from 2 mice (upper and lower row) in the naïve, learning and expert phases. Shaded bar depicts the stim period. Shadows are mean±SEM across trials (mouse 1: n=86/66, 90/70 and 166/173 Hit-Hit/CR-Hit for naïve, learning and expert phases respectively. mouse 6: n=94/80, 86/121 and 99/135) **b**. Grand average of BC activity during the stim period (−0.2:0.6ms) for the naïve, learning and expert phases. Boxes indicate quartiles at 25/75^th^ percentile across mice (n=7). **c**. Same as **a** but for body and whisker movements in the Hit-Hit (light gray) and CR-Hit (dark gray) trials. **d**. Same as **b** but for body (top) and whisker (bottom) movements. *p < 0.05; n.s. not significant; Wilcoxon signed-rank test.

We next quantified the emergence of history information with regard to the different time scales, the trial structure (within seconds) or the learning profile (across days). We first show 2D activity plots in BC for each trial pair (i.e., CR-Hit and Hit-Hit; showing activity of only the Hit trial), where trial time is plotted on the x-axis and trial number across learning time on the y-axis (Fig. 3a; 100-trial bins regardless of trial pair). Both trial pairs display an increase in activity during the stim-period shortly after passing the learning threshold. We defined a history modulation index as the difference in activity for BC between the two pair types (Hit-CR minus Hit-Hit). History modulation increased around the stim-period only in learning and expert phases but not in the naïve case (Fig 3b, c). A significant history modulation was defined as values exceeding mean±2SD of a trial-shuffled sample distribution (n=1000 iterations) and was performed for each mouse separately (Fig. 3b). The onset of the history modulation was defined as the first time frame reaching significant values (red arrows in Fig. 3b) and was found in BC to be during the stim-period (Fig. 3d; 0.05±0.32s, −0.1±0.27s 1s, median±SEM relative to texture stop in learning and expert phases respectively). We note that in the expert phase there is also a small peak exceeding the significance around the cue, indicating history information in BC may be present to some extent before stimulus presentation. Next, we quantified the history modulation in BC during the stim period as a function of the learning time course. History modulation in BC had the steepest increase after each mouse crossed its learning threshold (Fig. 3e, f). The onset of the history modulation was defined as the first trial bin exceeding the trial-shuffled sample distribution and was found to occur shortly after the learning threshold, highly correlated with the learning threshold indicating strong relationship between history emergence and learning of each individual mouse (Fig. 3g, h; 500±83 trials, median±SEM, r=0.97 p<0.001, spearman correlation). Note that our onset measurement is relatively strict and an increase in history information can be observed shortly (i.e., tens of trials) after crossing the learning threshold (Fig. 3d).

**Figure 3.**
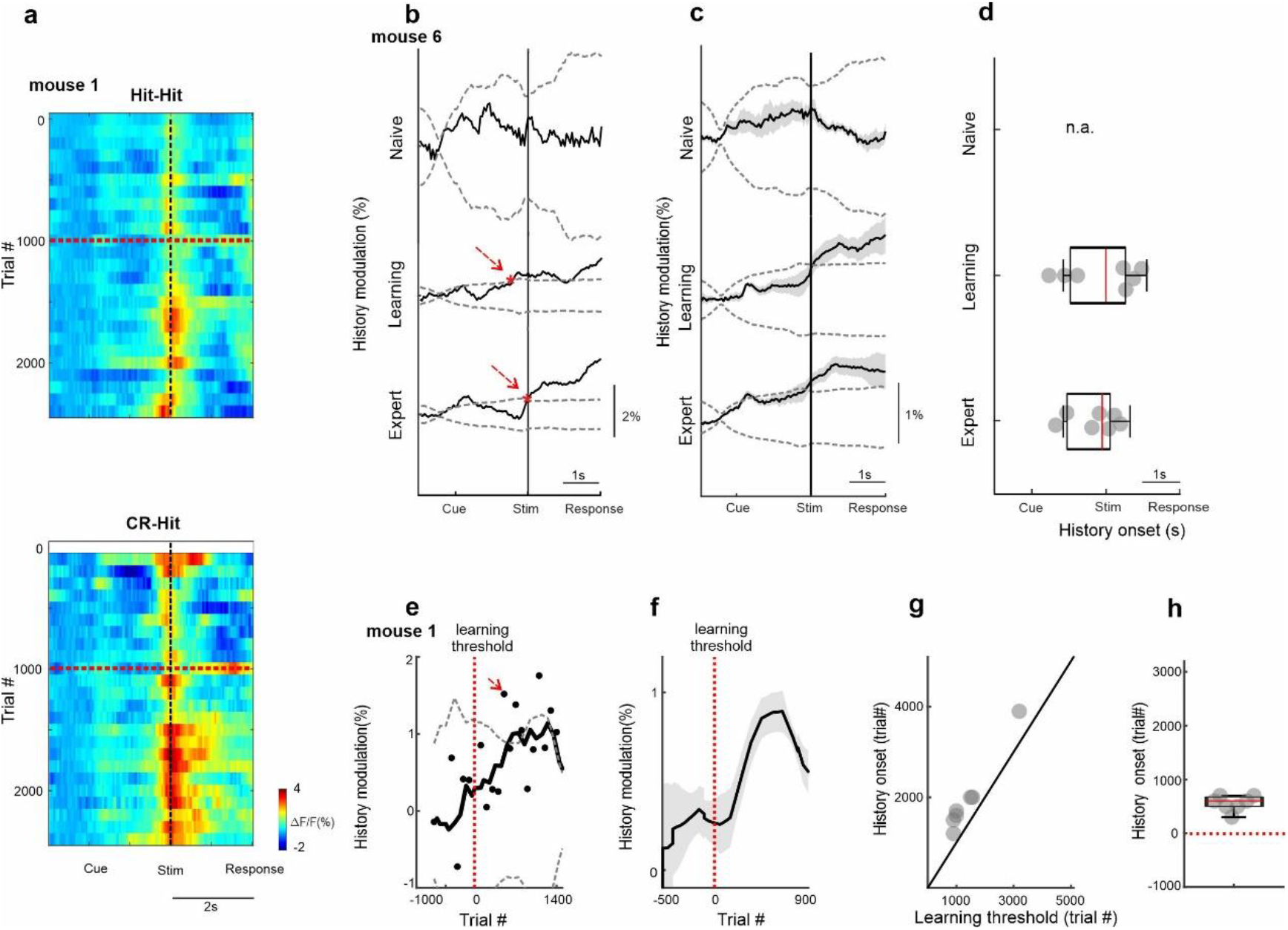
Temporal dynamics of history information in BC. **a**. 2D plot of BC responses for Hit-Hit (top) and CR-Hit (bottom; trial structure on x-axis; Trial number across learning (in bins of 100 trials) on the y-axis. Red horizontal dashed line indicates learning threshold. Black dashed vertical line indicates the time of texture stop. **b**. Example from one mouse of the history modulation (activity in CR-Hit minus activity in Hit-Hit) in BC along the trial structure in the naïve, learning and expert phases. Dashed gray line is the mean ± 2 SD of the trial-shuffled data (n=1000 iterations). The first-time frame crossing the shuffle data is defined as the onset and is marked in red. **c**. Mean history modulation in BC along trial time. Shadows depict mean±SEM across mice (n=7). **d**. Median onset of history modulation. Boxes indicate quartiles at 25/75^th^ percentile across mice (n=7). **e**. Example from one mouse of the history modulation along learning dimension. Dashed gray line is the mean ± 2 SD of the trial-shuffled data (n=1000 iterations). The first-time frame crossing the shuffle data is defined as the onset for learning and is marked in red. The vertical red dashed line (trial 0) marks the learning threshold. **f**. Mean history modulation in BC along the learning profile aligned to the learning threshold of each mouse (time 0). Shadows depict mean±SEM across mice (n=7). **g**. Onset of the history modulation for learning as a function of the learning threshold. Each point is one mouse (n=7). **h**. Median onset of history modulation relative to the learning threshold. Boxes indicate quartiles at 25/75^th^ percentile across mice (n=7).

We expanded our history analysis also for the pair types other than CR-Hit and Hit-Hit. For sufficient trial numbers, we focused on the learning phase. First, we compare FA-Hit to Hit-Hit and CR-Hit, i.e., the same current trial type but preceded by an error trial (FA). Response in BC for FA-Hit was similar to Hit-Hit and significantly lower compared to CR-Hit (Fig. s3; p<0.05 signed rank test). This result highlights that specifically a correct rejection (CR), rather than the stimulus (i.e., texture) type, has a strong history effect. Next, we compared FA-CR, Hit-CR and CR-CR, i.e., similar to the previous comparison differing only in the current trial type (CR instead of Hit). There was no significant difference between the different pairs, indicating that the current trial type, i.e., Hit in this case, has a strong effect along with the history of the CR (Fig. s3; p>0.05, signed rank test). A comparison of FA-FA, Hit-FA and CR-FA did not show a significant difference (Fig. s3; p>0.05, signed rank test). In general, a preceding CR trial resulted in higher activation independent of the current trial type (i.e., Hit, CR or FA; not significant for CR and FA), indicating that history information is present at the current time independently of incoming sensory information (Fig. s3; Compare orange bars to the blue bars). In conclusion, we found that the CR-Hit pair displayed a specific enhancement in BC that is related both to the preceding and current trial type (see discussion).

Next, we expanded our analysis to the whole dorsal cortex during the stim period. Mean activation maps for both CR-Hit and Hit-Hit pairs (i.e., Activity for the current Hit trial whereas only the preceding trial was different) during the stim period displayed a pronounced activation patch in BC during naïve, learning and expert phases (Fig. 4a). BC activity was higher in CR-Hit compared to Hit-Hit especially during learning and expert phases. The grand average activity for all 25 cortical areas highlights history-dependent information that emerges during learning (Fig. 4b). We note that other areas, e.g., different association areas, also encoded history-dependent information especially during learning and expert phases. Taken together, these results indicate that BC encodes history-dependent information that emerges during the stim period and just after learning. These results gave us the motivation to examine history-dependent information at time periods before texture touch.

**Figure 4.**
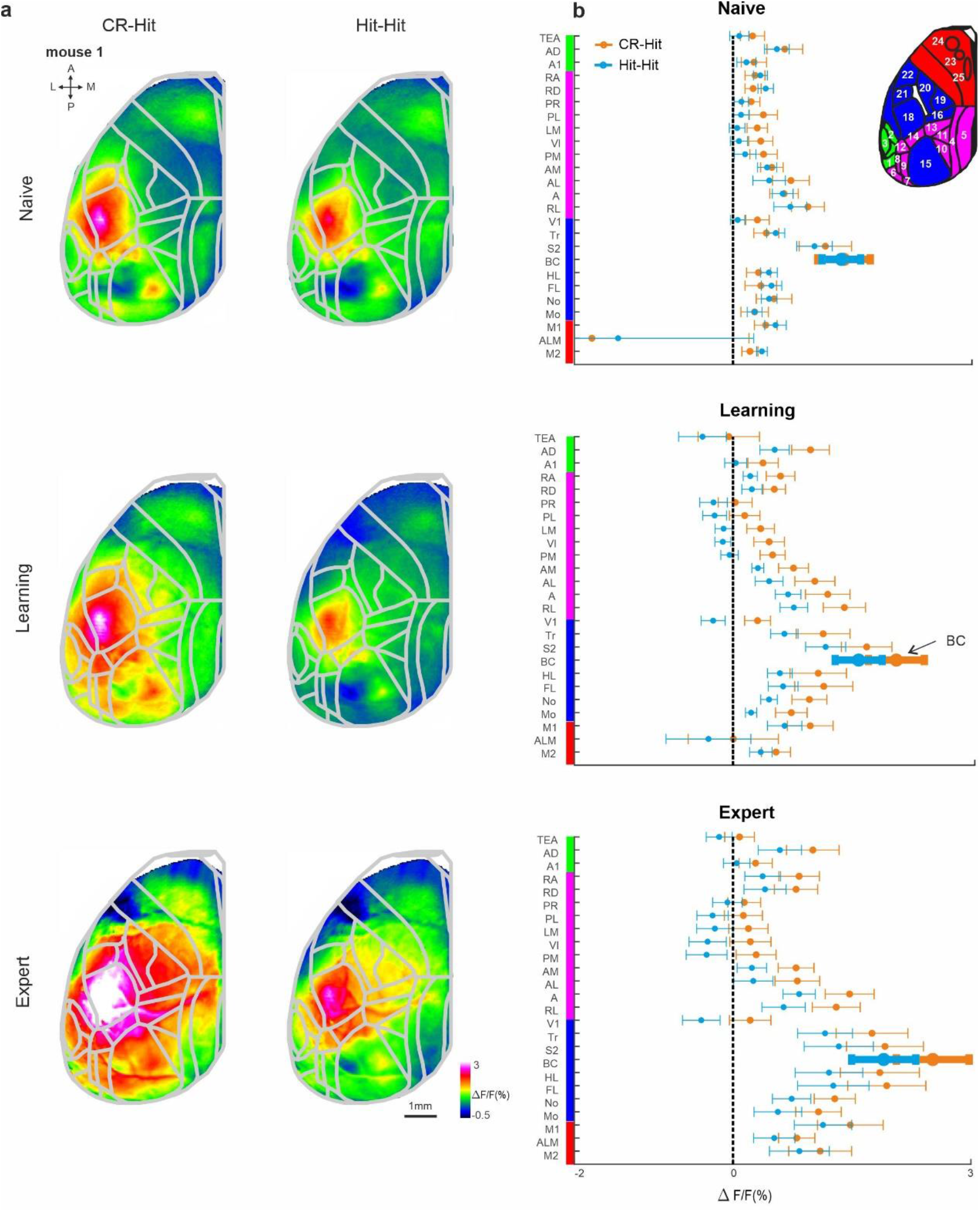
Cortex-wide history modulation during the stim period. **a**. Mean activity maps averaged within the stim period (−0.2 – 0 seconds relative to texture stop) of CR-Hit (left) Hit-Hit (right) during the naïve (top), learning (middle), expert (bottom) phases. Color bar denotes normalized fluorescence (ΔF/F). 2D top-view atlas is superimposed in gray. **b**. Grand average neuronal activity during the stim period (−0.2:0.2s) for Hit-Hit (blue) and CR-Hit (orange) in all 25 areas for the naïve (top), learning (middle) and expert (bottom) phases. Error bars depict mean±SEM across mice (n=7).

### History information in RL before sensation

We next focused our analysis on the pre-period, just before texture touch (−1 to −0.6 sec before texture stop). Mean activity maps during the pre-period highlight activity in association area rostrolateral (RL) that is present for both CR-Hit and Hit-Hit pairs during the naïve, learning and expert phases (Fig. 5a;(Gilad and Helmchen, 2020)) RL pre-period activity is higher in CR-Hit compared to Hit-Hit mostly during learning and expert phases. In addition, higher RL activity in CR-Hit pair starts even before the pre-period, indicating that history-information is not directly related on the current stimulus (Fig. 5b). The grand average of all 25 cortical areas, highlights the emergence of history-dependent information emerging during learning, especially in RL, but also in other association and sensory areas (Fig. 5c).

**Figure 5.**
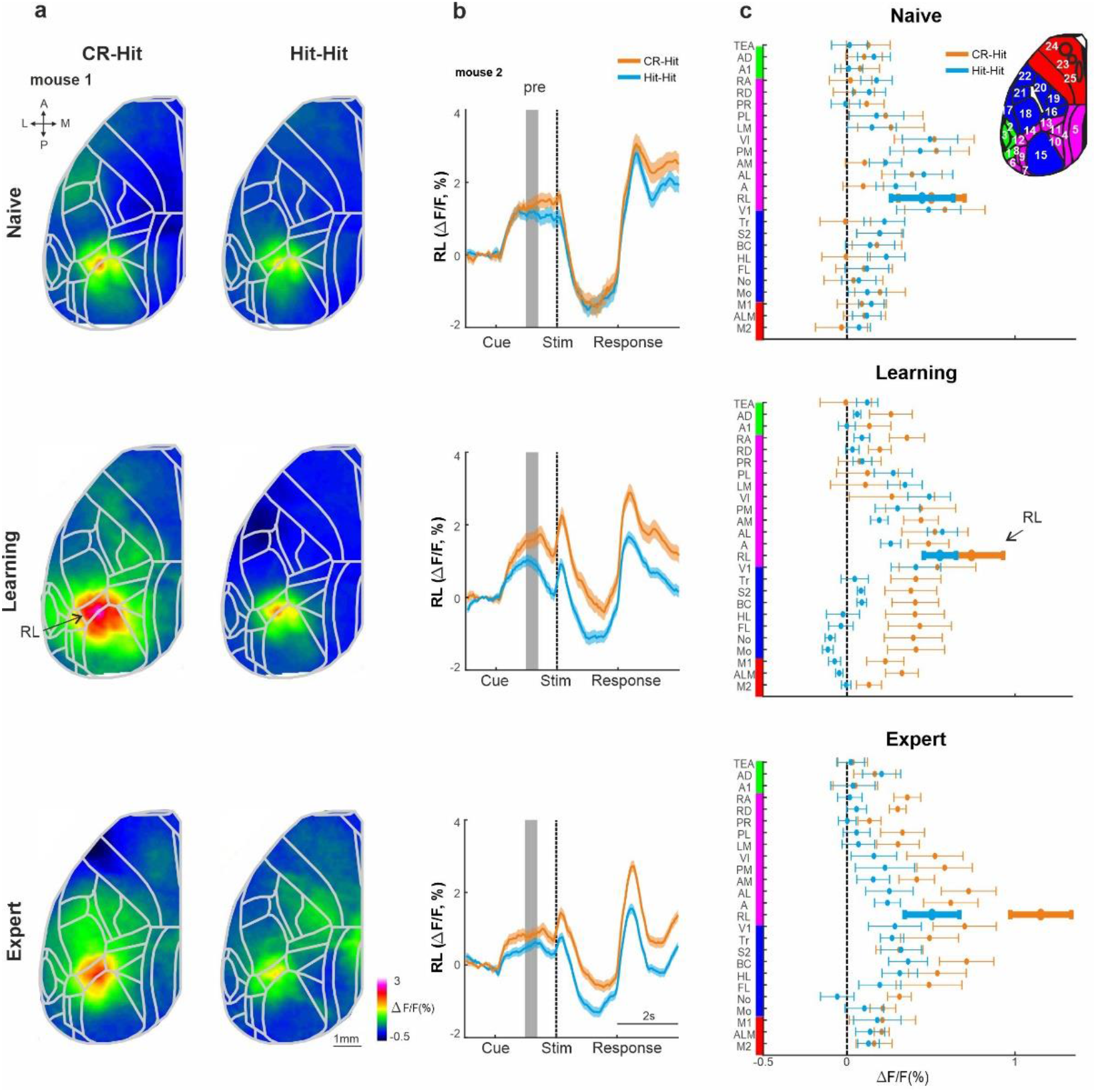
History information in RL before stimulus presentation. **a**. Mean activity maps averaged within the pre-period (−1 – −0.8 seconds relative to texture stop) of CR-Hit (left) Hit-Hit (right) during the naïve (top), learning (middle), expert (bottom) phases. Color bar denotes normalized fluorescence (ΔF/F). 2D top-view atlas is superimposed in gray. **b**. Example from one mouse of average RL response of Hit-Hit (blue) and CR-Hit (orange) in the naïve (top), learning (middle) and expert (bottom) phases. Shaded gray bar depicts the pre-period (−1– −0.6). Shadows are mean±SEM across trials (n=51/54, 92/78 and 168/173 Hit-Hit/CR-Hit for naïve, learning and expert phases respectively) **c**. Grand average neuronal activity during the pre-period (−1 – −0.6) for Hit-Hit (blue) and CR-Hit (orange) in all 25 areas for the naïve (top), learning (middle) and expert (bottom) phases. Error bars depict mean±SEM across mice (n=7).

RL activity was significantly higher in CR-Hit compared to Hit-Hit trials in the pre-period during the expert phase (Fig. s4; signed rank test, p<0.05, similar trend for the learning phase but insignificant; not significant for the naïve phase). The onset of history modulation within the trial structure (as in Fig. 3d) was earlier in RL compared to BC in both learning (−0.15±0.32s and 0.05±0.32s, median±SEM in RL and BC respectively) and expert phases (−0.75±0.2s and −0.1±0.27s, median±SEM in RL and BC respectively) but not significantly different (p>0.05, signed rank test). The onset for the history modulation with relation to the learning profile in RL (similar to Fig. 3h; During the pre-period) was also earlier than BC, but not significantly different (200±162 trials after crossing threshold compared to 500±83 in BC; median±SEM, p>0.05 singed rank test). Taken together, these results indicate that as mice gain expertise, RL encodes history information before the next stimulus occurs, which may inform through its projections to BC where history information then could be integrated with information of the current incoming texture.

### Past versus present discrimination power in BC and RL

How well can BC and RL activity discriminate at the single trial level history information compared to the information of the current stimulus? To do this, we computed the receiver operating characteristics (ROC) analysis between specific trial types(Gilad et al., 2020, 2013), along with the area under the curve (AUC) quantifying the discrimination power at the single trial level (Methods). We calculated the AUC between two types of trials (Fig. 6a): 1) Activity between CR-Hit and Hit-Hit pairs based on the activity during the Hit trial. This is defined as ‘history-AUC’ since only the previous trial is different. 2) Activity between the current Hit and CR trials. This is defined as the ‘Current-AUC’ because the current trial types are different (both in terms of stimulus type and action). Both history-AUC and current-AUC are calculated for BC and RL for each time frame along the trial structure and for naïve, learning and expert phases. Intuitively, one would assume that the current-AUC will display higher discrimination power compared to the history-AUC because the latter AUC measure compares the same current trial type which should be harder to discriminate. Interestingly, during the expert phase, history-AUC in both BC and RL has a discrimination power in the stim period that is not significantly different than that of the current-AUC (Fig. 6b, c; p>0.05; singed rank test). In other words, we found that BC and RL discriminate past stimuli just as well as the current stimuli. In addition, during the learning phase, RL and to some extent BC, display a significantly higher history-AUC compared to the current-AUC, specifically in the pre-period (Fig. 6d, e; p<0.05; Singed rank test). This indicates that history information is discriminative at the single trial level before stimulus onset. Taken together, we find that BC and RL can encode the past just as well as the present.

**Fig. 6.**
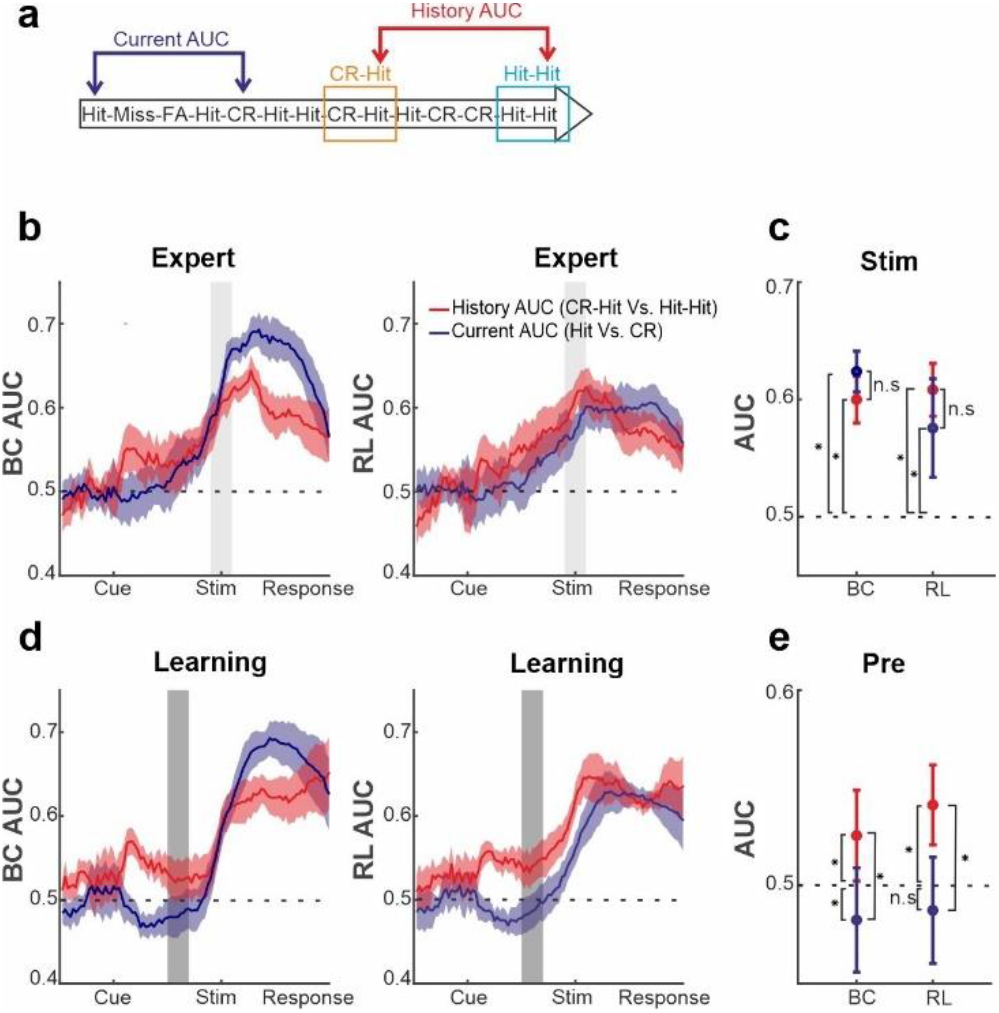
History and current information are equally discriminative at the single trial level. **a**. Schematic diagram for the two types of area under the curve (AUC) measures (derived from a ROC analysis): history-AUC between the Hit responses for Hit-Hit and CR-Hit trial types. Current-AUC between Hit and CR trial types regardless of their history. **b**. Grand average of the history (red) and current (blue) AUC measures in BC (left) and RL (right) along the trial structure during the expert phase. Shadows depict mean±SEM across mice (n=7). Values significantly differ from chance (0.5) in history-AUC (p<0.05, 2 tail ttest, for both BC and RL). **c**. Grand average of history and current-AUC measures during the stim period in the expert phase. Error bars indicate mean±SEM across mice(n=7). **d**. Same as in a but for the learning phase. Error bars as in a, values significantly differ from chance (0.5) for history-AUC (p<0.05, 2 tail ttest, for both BC and RL), but not for the current-AUC in RL. **e**. same as in c, but for the pre-period during the learning phase. *p < 0.05; n.s. not significant; Wilcoxon signed-rank test.

## Discussion

### History information is trial-type specific

We have identified cortex wide encoding of history information that emerges as mice learn to discriminate between two textures. History information was not dependent on the current stimulus and emerged in RL association area before texture touch. Our results indicate that a previous CR trial will lead to higher activity in BC and RL compared to a previous Hit trial. This difference is probably not due to pure sensory differences in the previous trial since the effect was not present after FA trials (sup Fig. s3, left panel). In addition, mice trained to lick the P1200 texture displayed a similar bias to the CR-Hit, further indicating that these differences are not purely sensory related. Moreover, this difference is probably not related to the previous motor action (e.g., either lick or no-lick). During the current trial, body and whisker movements were not significantly different, emphasizing that there are no motor-related differences based on the previous trial (Fig. 2c, d). The fact that these differences emerged only after learning implies that these differences are not purely sensory or motor related but rather reflect internal history-related information. It may be that in a go/no-go discrimination task the mouse mainly learns not to lick for the no-go texture (i.e., CR) making the information of a CR trials more pronounced relatively to a Hit trials. Another possibility is that a previous CR will cause a pronounced anticipatory state for the incoming texture, leading to enhanced cortical activity. Again, we did not find any consistent differences in motor movements based on the previous trials making this possibility less likely. In summary, our results indicate that history-dependent information emerges internally in cortex as mice learn to discriminate between two stimuli.

### History information emerges in RL and transferred to BC

BC is considered a lower-order sensory area but encodes not only lower-order stimulus features(Chen et al., 2013; Estebanez et al., 2012; Garion et al., 2014; Safaai et al., 2013) but also higher-order information such as choice and reward value(Chéreau et al., 2020; Rodgers et al., 2021; Zuo and Diamond, 2019). We additionally found that BC carries history information during the sensation period which is related to the previous trial several seconds back. The presence of history information in lower-order areas such as BC is interesting by itself, but also raises the question of where is its origin. Interestingly, we show that history information emerges in RL before texture touch implying that RL may transfer history information in a top-down manner to BC for optimal sensory integration.

The presence of history information in RL before the sensation period implies that RL may play a crucial role in linking past experience to ongoing sensory integration. RL is the lateral part of PPC adjacent to BC, within the cluster of higher-order association areas surrounding V1 (Hovde et al., 2018; Lyamzin and Benucci, 2019). Previous studies showed that history information of choice-outcome is encoded by PPC neurons(Harvey et al., 2012; Hwang et al., 2017; Marcos and Harvey, 2016; Pho et al., 2018), as well as history of sensory information(Akrami et al., 2018). Silencing the PPC specifically during the inter-trial interval affected the behavioral performance of rats (Akrami et al., 2018; Hwang et al., 2017), whereas silencing during the stimulus presentation did not affected performance. The PPC is also reciprocally connected to hippocampus via entorhinal and retrosplenial cortices (Save and Poucet, 2009; Whitlock et al., 2008) and to basolateral amygdala via the anterior cingulate cortex (Suzuki et al., 2022), giving fast access to the different memory hubs. Khodagholy et al.(Khodagholy et al., 2017) showed coupling of PPC and hippocampal ripples that strengthen in non-REM sleep after rats learned a spatial exploration task, further indicating that RL may relay history information from subcortical memory hubs to cortex.

The fact that history information emerges only after learning, implies that it encodes a subjective value or association of a certain past stimulus. It may be that only once the value of a certain stimulus has been established, e.g., by strengthening indirect connections between basolateral amygdala (that has a role in associative memory) and RL, history information can aid in efficiently encoding the incoming stimulus. In light of this discussion, we suggest that the consolidation of a certain association (in our case a CR), induces long-term synaptic plasticity of top-down projections from higher-order association area (e.g., RL) to a lower-order sensory area (e.g., BC). This projection-specific potentiation may better recruit sensory cortex in the context of the immediate previous history.

### Mechanisms for integrating past and present

The wide-field signal measured in our study reports bulk population activity specifically in L2/3 excitatory cells. Are neuronal populations encoding past and present information in the BC overlapping or distinct? On one side, it could be that the same cell in BC encodes both the current stimulus and additionally receives top-down input from RL carrying the past stimulus identity. This additional top down information may amplify sensory integration and optimize discrimination of the current stimulus. On the other side, previous studies that measured single cell activity in the BC showed that single cells tend to respond to one information type, (Chéreau et al., 2020; Estebanez et al., 2012; Rodgers et al., 2021). In this case, we hypothesize that different populations in BC encode current and history information, which leads to a larger fraction of neurons in BC that are active for the CR-Hit pair. A larger number of active neurons in BC may facilitate sensorimotor integration involving downstream areas such as the motor cortex, further resulting in gaining an expert level (Zuo and Diamond, 2019).

It is probable that both history and learning involve other circuit elements such as deep cortical layers (Pasupathy and Miller, 2005; Roelfsema and Holtmaat, 2018; Vecchia et al., 2020), inhibitory subtypes, other pathways (Lacefield et al., 2019; Mohan et al., 2022; Musall et al., n.d.; Petreanu et al., 2012; Williams and Holtmaat, 2019),and subcortical areas (Fu et al., n.d.; Garrett et al., 2020; Pasupathy and Miller, 2005; Pfeffer et al., 2013). Future work may aim to dissect specific subpopulations that carry history information using similar behavioral tasks, e.g., imaging of cortex-wide layer 5 dynamics. Layer 5 neurons may be ideal in integrating past information arriving onto the apical dendrites in layer 1^54^ with incoming information arriving from the thalamus. In addition, similar tasks with reward after CR trials, or tasks that better differentiate between choice and outcome (decision tasks, giving different probabilities of outcome to each choice), or tasks with a dynamic inter-trial interval may shed light on the meaning of this history-learning effect. In summary, our results imply that as we learn, the cortex learns to better integrate past and present information resulting in expert performance.

## Acknowledgements

This project has received funding from the European Union’s Horizon 2020 research and innovation program under the Marie Skłodowska-Curie grant agreement No 659719 (A.G.). This work was supported by grants from Hebrew University of Jerusalem (Start-up grant; A.G.) and the Swiss National Science Foundation (SNSF) (31003A-149858 and 310030B_170269; F.H.).

## Data and code availability

The data and code that support the findings of this study are available at https://osf.io/hkvc5.

## Author contributions

A.G. and F.H. designed the experiments. A.G. conducted the experiments. A.G. and O.M. performed data analysis. A.G. and O.M. wrote the manuscript with comments from F.H.

## Star methods

### Animals and surgical procedures

Methods were carried out according to the guidelines of the Veterinary Office of Switzerland and following approval by the Cantonal Veterinary Office in Zurich. A total of 7 adult male mice (1-4 months old) were used in this study. These mice were triple transgenic Rasgrf2-2A-dCre; CamK2a-tTA;TITL-GCaMP6f animals, expressing GCaMP6f in excitatory neocortical layer 2/3 neurons (Gilad and Helmchen, 2020). The dataset used here is identical to our previous study (Gilad and Helmchen, 2020), but here we have applied a completely novel history analysis. To generate triple transgenic animals, double transgenic mice carrying CamK2a-Tta62 and TITL-GCaMP6f63 were crossed with a Rasgrf2-2A-dCre line (64; individual lines are available from The Jackson Laboratory as JAX# 016198, JAX#024103, and JAX# 22864, respectively). The Rasgrf2-2A-dCre;CamK2a-tTA;TITL-GCaMP6f line contains a tet-off system, by which transgene expression can be suppressed upon doxycycline treatment ((Garner et al., 2012; Gossen and Bujard, 1992). However, doxycycline treatment is not necessary in these animals, since the Rasgrf2-2A-dCre line holds an inducible system of its own, given that the destabilized Cre (dCre) expressed under the control of the Rasgrf2-2A promoter needs to be stabilized by trimethoprim (TMP) to be fully functional. TMP (Sigma T7883) was reconstituted in Dimethyl sulfoxide (DMSO, Sigma 34869) at a saturation level of 100 mg/ml, freshly prepared for each experiment. For TMP induction, mice were given a single intraperitoneal injection (150 μg TMP/g body weight; 29 g needle; 3–5 days post-surgery), diluted in 0.9% saline solution. We used an intact skull preparation (Silasi et al., 2016) for chronic wide-field calcium imaging of neocortical activity(Gilad et al., 2018b). Mice were anesthetized with 2% isoflurane (in pure O2) and body temperature was maintained at 37 °C. We applied local analgesia (lidocaine 1%), exposed and cleaned the skull, and removed some muscles to access the entire dorsal surface of the left hemisphere (Fig. 2a; ~6 × 8 mm2 from ~3 mm anterior to bregma to ~1 mm posterior to lambda; from the midline to at least 5 mm laterally). We built a wall around the hemisphere with adhesive material (iBond; UV-cured) and dental cement “worms” (Charisma). Then, we applied transparent dental cement homogenously over the imaging field (Tetric EvoFlow T1). Finally, a metal post for head fixation was glued on the back of the right hemisphere. This minimally invasive preparation enabled high-quality chronic imaging with high success rate.

### Texture discrimination task

Mice were trained on a go/no-go discrimination task (Fig. 1a) using a data acquisition interface (USB-6008; National Instruments) and custom-written LabVIEW software (National Instruments). Each trial started with an auditory cue (stimulus cue; 2 beeps at 2 kHz, 100-ms duration with 50-ms interval), signaling the approach of either two types of sandpapers (grit size P100: rough texture; P1200: smooth texture; 3M) to the mouse’s whiskers as ‘go’ or ‘no-go’ textures (Fig. 1a; pseudo-randomly presented with no more than three repetitions). Sandpapers were mounted onto panels attached to a stepper motor (T-NM17A04; Zaber) mounted onto a motorized linear stage (T-LSM100A; Zaber) to move textures in and out of reach of whiskers. The texture stayed in touch with the whiskers for 2 s, and then it was moved out after which an additional auditory cue (response cue; 4 beeps at 4 kHz, 50-ms duration with 25-ms interval) signaled the start of a 2-s response period. The stimulus and response cues were identical in both textures. The interval between the trails was 6 s (8 s from response to next cue). A water reward (~3 μL) was given to the mouse for licking for the go texture only after the response cue (‘Hit’), i.e. for the first correct lick during the response period (Fig. 1a; lick were detected using a piezo sensor). Punishment with white noise was given for licking for the no-go texture (‘false alarms’; FA). Licking before the response cue was neither rewarded nor punished. Reward and punishment were omitted when mice withheld licking for the no-go (‘correct-rejections’, CR) or go (‘Misses’) textures.

#### Training and performance

Five mice were trained to lick for the P100 texture (mice #1-4 and 6) and 2 mice were trained to lick for the P1200 texture (mice #5 and 7). Mice were first handled and accustomed to head fixation before starting water scheduling. Before imaging began mice were conditioned to lick for reward after the go texture (presented within a similar trial structure as the task itself). Imaging began only after mice reliably licked for the response cue (typically after the first day; 200–400 trials). On the first day of imaging, mice were presented with the ‘go’ texture and after 50 trials the ‘no-go’ texture was gradually introduced (starting from 10% and increasing by 10% approximately every 50 trials (Guo et al., 2014) until reaching 50% probability for the no-go texture by the end of the day. 6 out of the 7 mice learned the task within 3–4 days after around a thousand trials (Supplementary Fig. 1). Mouse #4 learned the task within 10 days. An effort was made to maintain a constant position of the texture and cameras across imaging days in order to maintain similar stimulation and imaging parameters.

#### Wide-field calcium imaging

We used a wide-field approach to image large parts of the dorsal cortex while mice learned to perform the task (Gilad et al., 2018b) .A sensitive CMOS camera (Hamamatsu Orca Flash 4.0) was mounted on top of a dual objective setup. Two objectives (Navitar; top objective: D-5095, 50 mm f0.95; bottom objective inverted: D-2595, 25 mm f0.95) were interfaced with a dichroic (510 nm; AHF; Beamsplitter T510LPXRXT) filter cube (Thorlabs). This combination allowed a ~9-mm field-of-view, covering most of the dorsal cortex of the hemisphere contralateral to texture presentation. Blue LED light (Thorlabs; M470L3) was guided through an excitation filter (480/40 nm BrightLine HC), a diffuser, collimated, reflected from the dichroic mirror, and focused through the bottom objective ~100 μm below the blood vessels. Green light emitted from the preparation passed through both objectives and an emission filter (514/30 nm BrightLine HC) before reaching the camera. The total power of blue light on the preparation was <5mW; i.e., <0.1 mW/mm2. At this illumination power we did not observe any photobleaching. Data was collected with a temporal resolution of 20 Hz and a spatial sampling of 512 × 512 pixels, resulting in a spatial resolution of ~20 μm/pixel. On each imaging day a green reflectance image was taken as reference to enable registration across different imaging days using the blood vessel pattern (fibercoupled LED illuminated from the side; Thorlabs).

#### Mapping and area selection

Each mouse underwent a mapping session under anesthesia (1% isoflurane), in which we presented five different sensory stimuli (contra-lateral side (Gilad and Helmchen, 2020). Next, we registered each imaging day to the mapping day using skull coordinates from the green images. Finally, we registered each mouse onto a 2D top view mouse atlas using both functional patches from the mapping and skull coordinates ((Gilad and Helmchen, 2020);©2004 Allen Institute for Brain Science. Allen Mouse Brain Atlas. Available from: http://mouse.brain-map.org/29). Within the atlas borders, we defined 25 areas of interest, with some manual modifications within these borders to fit the functional activity for each mouse. Motor cortex areas were defined based on stereotaxic coordinates and functional patches for each mouse (see below). Thus, all mice had similar regions of interest that were comparable within and across mice. We grouped these 25 areas into auditory (green), association (pink), somatosensory + V1 (blue), and motor (red) areas (Fig. 1d and Supplementary Fig. 1b). Auditory areas: Primary auditory (A1), Auditory dorsal (AD) and Temporal association area (TEA). Sensory areas: Somatosensory mouth (Mo), Somatosensory nose (No), Somtosensory hindlimb (HL), Somtosensory forelimb (FL), Barrel cortex (BC; Primary somatosensory whisker); Secondary somatosensory whisker (S2), Somtosensory trunk (Tr) and Primary visual cortex (V1). Motor areas: whisker-related primary motor cortex (M1; 1.5 anterior and 1mm lateral from bregma, corresponding to the whisker evoked activation patch in M1 from the mapping session), anterior lateral motor cortex (ALM; 2.5 anterior and 1.5 mm lateral from bregma69) and secondary motor cortex (M2; 1.5 anterior and 0.5mm lateral from bregma corresponding11). Association cortex: Rostrolateral (RL), Anterior (A), Anterior lateral (AL), Anterior medial (AM), Posterior medial (PM), Lateral medial (LM), Lateral intermediate (LI), Posterior lateral (PL), Post-rhinal (PR), Retrosplenial dorsal (RD) and Retrosplenial angular (RA). We note that our definition of association cortex is broad and may include or exclude areas that are not necessarily classical association areas.

#### Whisker and body tracking

In addition to wide-field imaging, we tracked movements of the whiskers and the body of the mouse during the task (Fig. 1a). The mouse was illuminated with a 940-nm infrared LED. Whiskers were imaged at 50 Hz (500 × 500 pixels) using a high-speed CMOS camera (A504k; Basler), from which we calculated time course of whisking envelope and the time of first touch (see below). An additional camera monitored the movements of the mouse at 30 Hz (The imaging source; DMK 22BUC03; 720 × 480 pixels). We used movements of both forelimbs and the head/neck region to assess body movements, to reliably detect large movements (Fig. 1a; see Data Analysis below).

#### Calculating body movements

We used a body camera to detect general movements of the mouse (30 Hz frame rate). For each imaging day, we first outlined the forelimbs and the neck areas (one area of interest for each), which were reliable areas to detect general movements. Next, we calculated the body movement (1 minus frame-to-frame correlation) within these areas as a function of time for each trial. We than averaged all the defined body areas to one “body” vector.

#### Whisker tracking

The average whisker angle across all imaged whiskers was measured using automated whisker tracking software (Knutsen et al., 2004). The mean whisker envelope was calculated as the difference between maximum and minimum whisker angles along a sliding window equal to the imaging frame duration (50 ms; (Gilad et al., 2018b)). Whisker envelope was normalized just before the auditory cue similar to wide-field data (Frame zero). In addition, we manually detected the first frame, in which any whisker touched the upcoming texture, using the movies from the whisker camera (LabVIEW custom program). The first touch occurred on average 0.33 and 0.34 s before the texture stopped for naïve and expert mice respectively. Time of first touch did not differ between expert and naïve mice (P > 0.05; Mann–Whitney U-test; n = 7 mice). We note that the first touch occurred mostly (but not exclusively) in the pre-period from −1 to −0.5 relative to texture stop.

#### Data analysis

Data analysis was performed using Matlab software (Mathworks). All mice were continuously imaged during learning (5–11 days). Wide-field fluorescence images were sampled down to 256 × 256 pixels and pixels outside the imaging area were discarded. This resulted in a spatial resolution of ~40 μm/pixel and was sufficient to determine cortical borders, despite further scattering of emitted light through the tissue and skull. Each pixel and each trial were normalized to baseline several frames before the stimulus cue (frame 0 division). Our main focus was on the history effect. Because the hit trails had the largest portion from all trails, we focused on the hit trials. We sub grouped all the Hit trials based on the type of the preceding trial as follow: CR-Hit - Hit trials that were preceded by correct rejection trial. Hit-Hit - Hit trials that were preceded by a Hit trial. FA-Hit - hit trials that were preceded by a false alarm trial. We mainly focused on comparing Hit-Hit and CR-Hit pairs since they had a large proportion in naïve, learning and expert phases (but see Figure s4). We defined two time periods within the trial structure: pre (−1 to 0.6 s relative to texture stop) and stim (−0.2 to 0.2 relative to texture stop; Fig. 1a).

#### Calculation of learning curves

Trials were binned (n = 100 trials with no overlap) across learning (at the stimulus time, adjusted for each mouse) and the performance (defined as d′ = Z(Hit/(Hit +Miss)) – Z(FA/(FA + CR)) where Z denotes the inverse of the cumulative distribution function) was calculated for each bin. Next, each behavioral learning curve was fitted with a sigmoid function 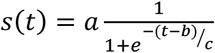 Where *a* denotes the amplitude, *b* the time point (in trial numbers) of the inflection point, and *c* the steepness of the sigmoid.

A learning threshold was defined as the bin in which the d’ crossed the inflection point (half point) of the learning curve sigmoid fit. (Fig. s1).

#### Defining the learning phases

We defined the naïve, learning and expert phase each as one day of recordings, the naïve day was defined as the first day to have enough correct rejections that the performance is still before the crossing threshold (typically the 2nd recording day). The learning day was defined as the day that the mouse crossed the learning threshold, and the expert was defined as the last day of the mouse (usually the 5th day)

#### Calculating history modulation and onset

We defined the ‘history modulation’ as the difference between the average activation of all CR-Hit and Hit-Hit trials. To calculate significance of history modulation, we calculated the sample distribution by trial shuffling between CR-Hit and Hit-Hit trials (n=1000 iterations). We than defined the onset of the history modulation as the first bin exceeding mean ± 2 SD of the sample distribution. We calculated this history modulation and significance across the trial dimension (every frame) and across learning dimension (every 100 trials). In the learning dimension, we calculated the average activity in the stim period (−0.2:0.2) of all the CR-Hit and Hit-Hit trials that were falling within each 100 trials bin.

#### Discrimination power between hit trials sub grouped by history

To measure how well could neuronal populations discriminate between go and no-go textures, we calculated a receiver operating characteristics (ROC) curve and calculated its area under the curve (AUC; with a value of 0.5 indicating no discrimination power). This can be done for a given area, each time frame within each learning phase separately (Fig. 6).

#### Statistical analysis

In general, the Wilcoxon signed-rank test to compare a population’s median to zero (or between two paired populations). Multiple group correction was used when comparing between more than two groups.

## Supplemental information

**Figure s1.**
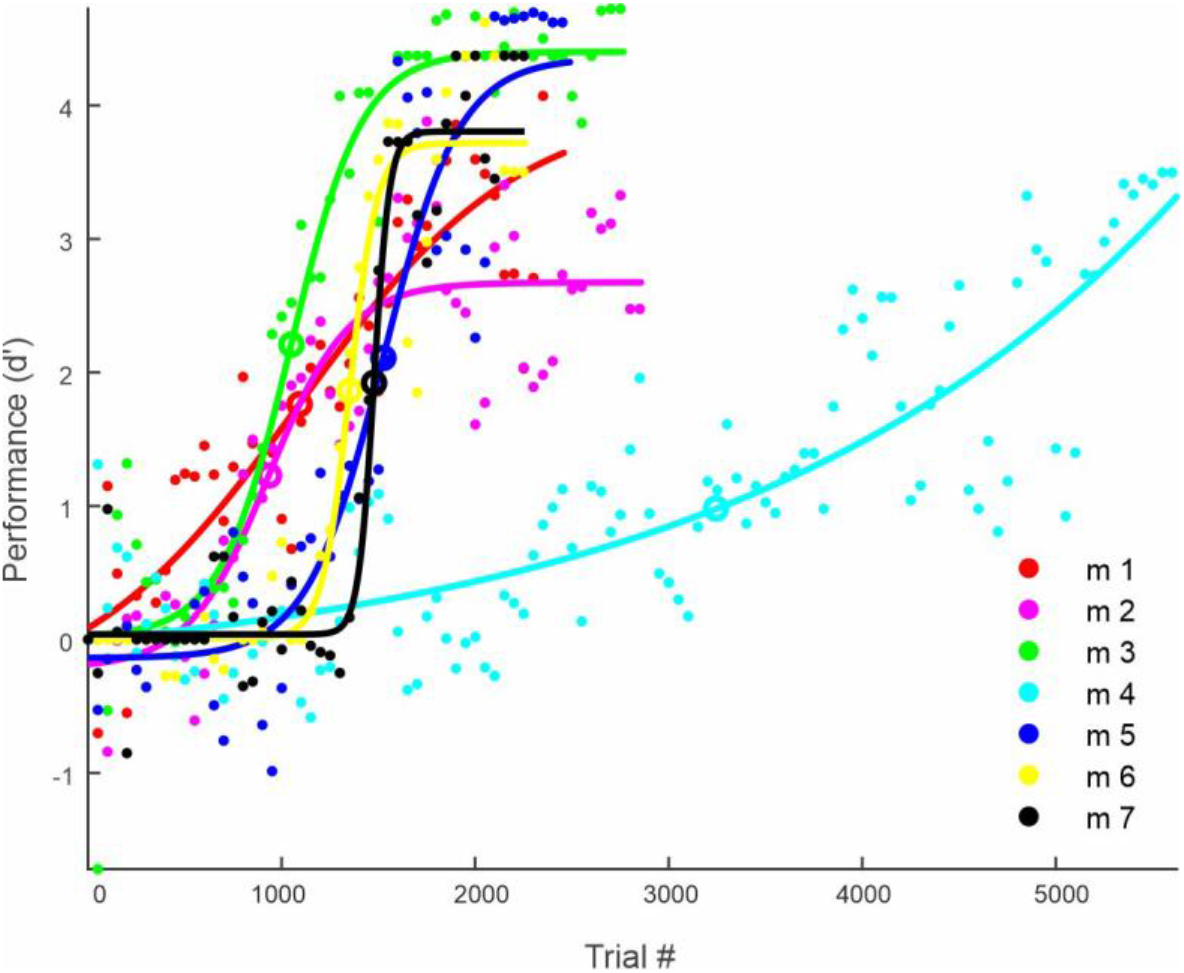
Learning curves of all 7 mice. Performance (d’) for all mice across the entire learning period is calculated for every 50 trails, fitted with a sigmoid function. The inflection point of the sigmoid fit is defined as the learning threshold and indicated by open circle for each mouse.

**Figure s2.**
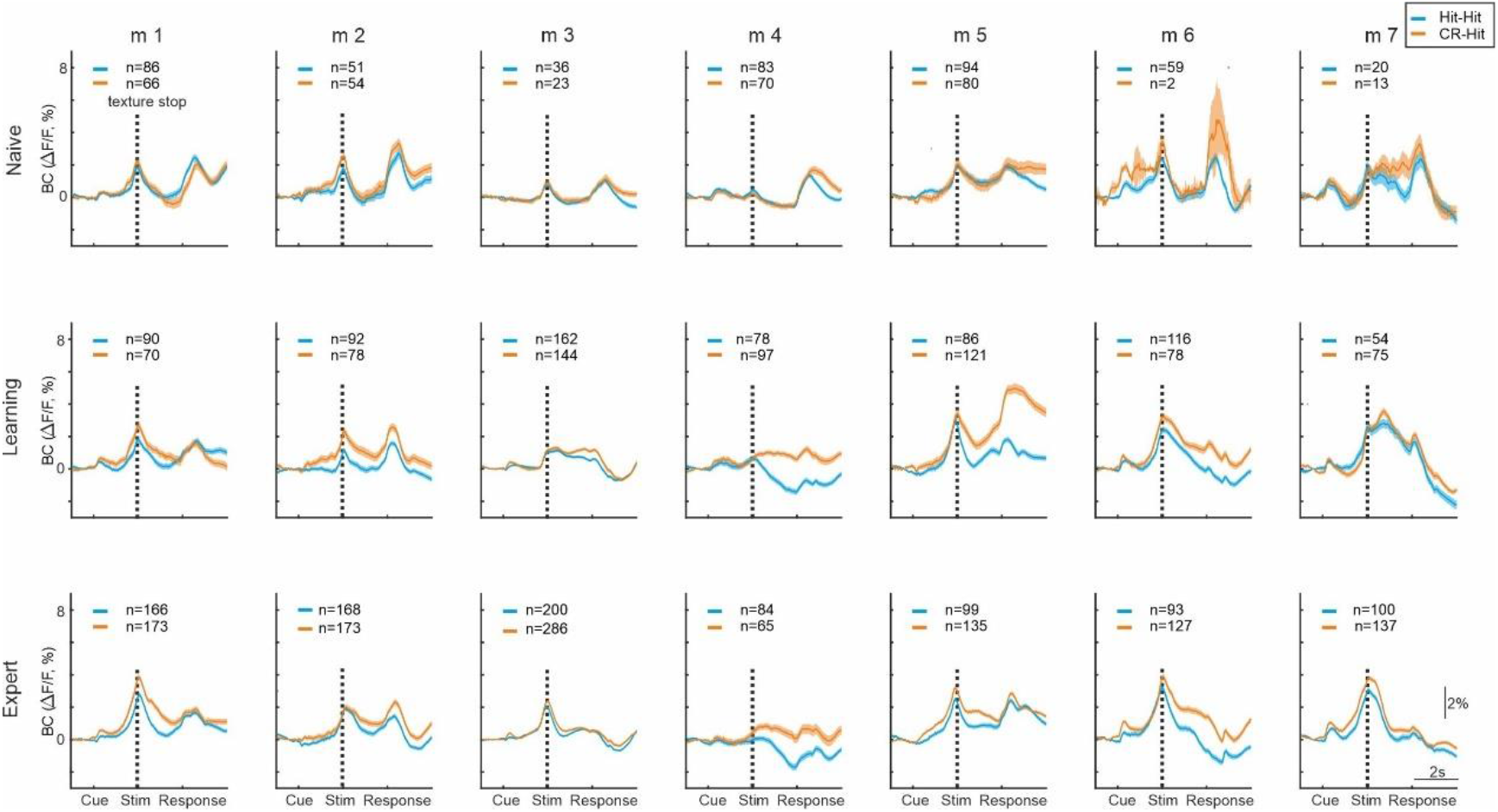
Time course of each mouse in the naïve, learning and expert phase for hit trials classified by preceding trial. Vertical dashed line denote texture stop. Shadows are mean±SEM across trials

**Figure s3.**
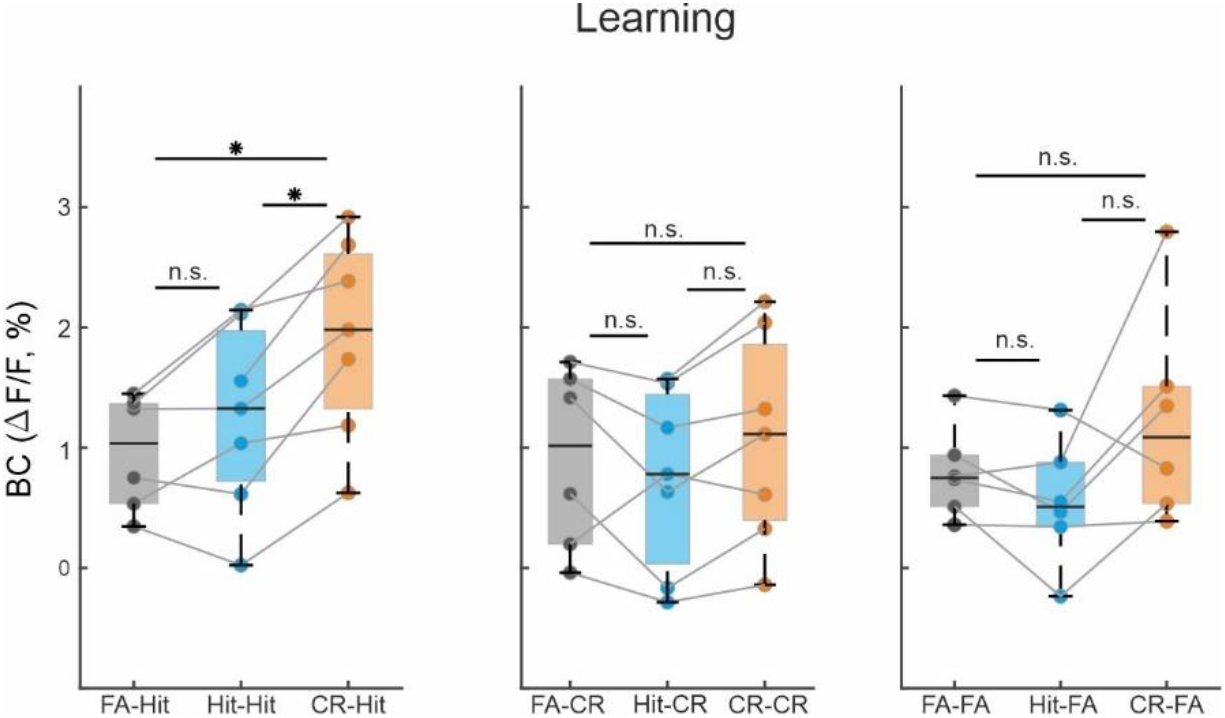
Correct rejection trials have the strongest history effect. Grand average of BC activity of all history combinations in the learning phase during the stim period (−0.2–0.6ms). Boxes indicate quartiles at 25/75^th^ percentile across mice (n=7). *p < 0.05; n.s. not significant; Wilcoxon signed-rank test.

**Figure s4.**
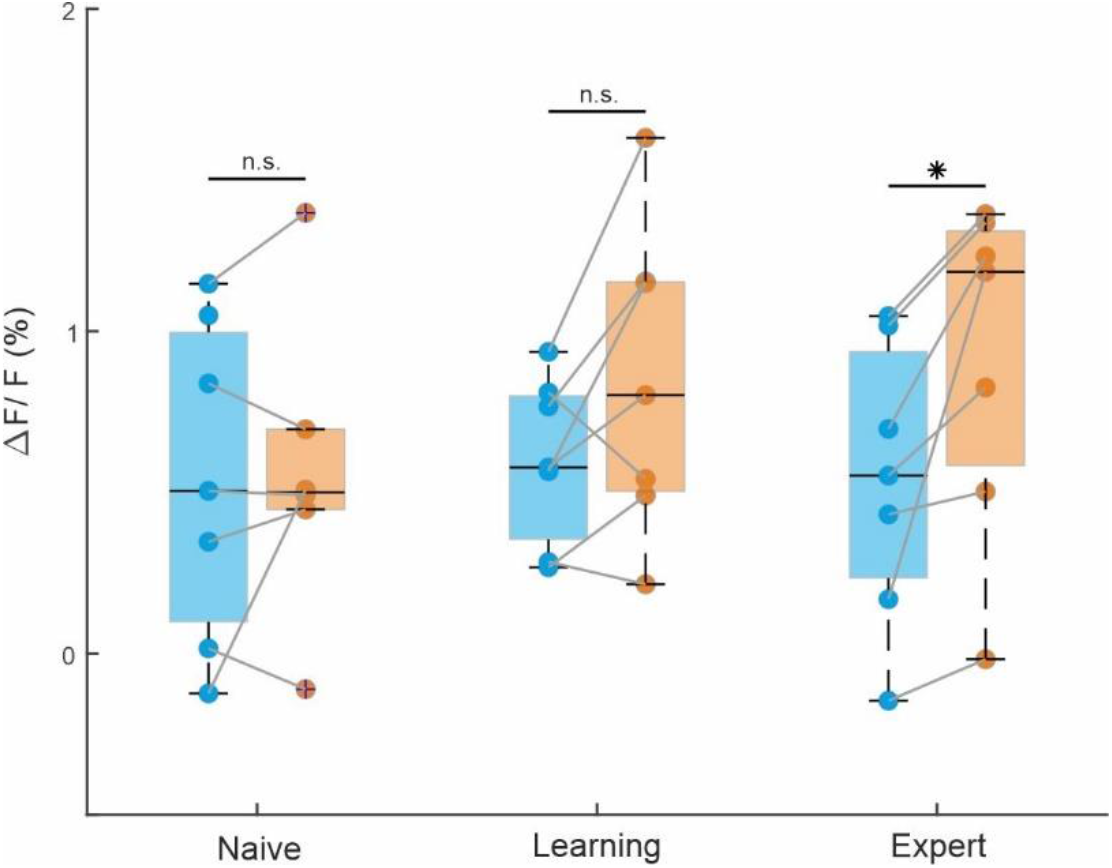
RL activity at pre-stim period (−1– −0.6) across learning. Boxes indicate quartiles at 25/75^th^ percentile across mice (n=7). *p < 0.05; n.s. not significant; Wilcoxon signed-rank test.

